# Apple floral attractants function across the 24-hour cycle to support potential diurnal and nocturnal pollinators

**DOI:** 10.64898/2026.11.13.766585

**Authors:** Sushant Potdar, Dmitry Kutcherov, Madison Jennings, Emily Brown, Conor Moriarty, My Tra Ly, Grace Hirzel, Tristan Hoerschelmann, Ngoc T. Phan, Neelendra Joshi, Erica L. Westerman

**Affiliations:** Department of Biological Sciences, University of Arkansas, Fayetteville, AR 72701, USA; Department of Molecular Biology & Genetics, Cornell University, Ithaca, NY 14853, USA; Department of Entomology, University of Arkansas, Fayetteville, AR 72701, USA; Department of Biological Sciences, North Dakota State University, Fargo, ND 58102, USA; Research Center for Tropical Bees and Beekeeping, Gia Lam, Hanoi 100000, Vietnam

**Keywords:** circadian rhythms, community ecology, floral volatiles, nectar, nocturnality, plant-insect interactions, pollination

## Abstract

Pollination in apple (*Malus domestica* Borkh.) is typically attributed to diurnal bees, yet nocturnal insects also contribute to fruit set. To assess whether apple flowers recruit potential pollinators throughout the 24-hour cycle, we quantified diel patterns in floral attractants and insect visitation in an orchard in Northwest Arkansas, United States. We measured nectar and volatiles and used a combination of visual surveys, light traps, and video recordings to assess insect abundance and floral visitation while controlling for weather conditions. We found that nectar volume remained consistent across sampling times, indicating that reward availability is dielly stable. Floral volatile bouquets, in contrast, exhibited subtle but significant diel and interannual variation, driven largely by shifts in aldehydes, alkanes, and benzenoids. Several compounds associated with nocturnal moth and dipteran attraction – including alpha-farnesene, benzyl alcohol, and aldehydes – were enriched at night. Apple flowers were visited by insects around the clock. Nighttime visitors were less abundant than daytime visitors but taxonomically diverse, including moths and nematoceran flies, whereas bees, wasps, and brachyceran flies were restricted to daytime. Different taxonomic groups therefore alternated over the diel cycle, producing pronounced shifts in visitor community composition. Thus, apple flowers produce attractants across the diel cycle and nocturnal insects – particularly dipterans and moths – are a consistent, ecologically meaningful component of the apple flower-visiting assemblage. Our findings support the need for pollinator-inclusive management that accounts for the full diel cycle, particularly when scheduling pesticide applications near bloom, to sustain diverse pollinator communities and enhance the resilience of apple production systems.

## 1. Introduction

Pollination is a fundamental ecological process sustaining both natural ecosystems and agricultural productivity. Animal pollinators – particularly insects – facilitate reproduction in the majority of flowering plants, thereby improving biodiversity, ecosystem stability, and global human food security (Klein et al. 2006, Potts et al. 2016, Pardo and Borges 2020, Gazzea et al. 2023). However, the widespread decline of managed and wild pollinator populations in recent decades has raised critical concerns for the sustainability of pollinator-dependent crops (Gallai et al. 2009, Asare et al. 2017), including apples (Pérez-Méndez et al. 2020). Traditionally, different species of bees have been used as the main pollinator for food production systems worldwide. The pollinator decline has compelled growers to rely increasingly on managed honey bee colonies, an expensive and ecologically limited solution that places additional stress on these insects and neglects the roles of native pollinators (Klein et al. 2006, Garibaldi et al. 2013, 2016, Olhnuud et al. 2022, Eeraerts et al. 2025). Identifying and supporting alternative pollination systems is therefore essential to maintaining resilient agricultural production.

Although bees are widely recognized as the primary pollinators in agricultural landscapes, they represent only a fraction of the diverse assemblage of insects that transport pollen. Numerous non-bee pollinators – including flies, beetles, and moths – contribute significantly to pollination services but remain understudied and underappreciated (Orford et al. 2015, Hahn and Brühl 2016, Sayers et al. 2019). Among these, moths and other nocturnal insects have received particularly little attention. Traditionally regarded as pests, nocturnal Lepidoptera have now been shown to function as efficient pollinators in a range of plant systems (Barthelmess et al. 2006, Amorim et al. 2013, Walton et al. 2020). Recent studies demonstrate that nocturnal insects can make substantial contributions to fruit and seed production, sometimes equaling or complementing the effects of diurnal pollinators (Cutler et al. 2012, Buxton et al. 2022, Fijen et al. 2023, Blareau and Requier 2025). Given their abundance, broad distribution, and distinct temporal activity patterns, nocturnal pollinators may represent a crucial yet overlooked component of pollination networks, especially as global bee populations continue to decline.

Apple (*Malus domestica* Borkh.) is among the most economically important fruit crops worldwide, contributing tens of billions of dollars annually to the global market (O’Rourke 2021). Apples are self-incompatible, meaning they require cross-pollination to produce a full crop, with effective insect-mediated pollination also contributing to fruit size and quality (McGregor 1976, Olhnuud et al. 2022). Unlike plants that exhibit nyctinasty, apple flowers stay open continuously day and night, remaining functional for an average of about seven days depending on the weather (Pratt 1988, Losada and Herrero 2013, Monteiro et al. 2015). Previous work in northwest Arkansas, United States, revealed that nocturnal insects – particularly moths – frequently visit apple blossoms and contribute substantially to seed set (Robertson et al. 2021). Moreover, flowers exposed to both diurnal and nocturnal visitors produced yields similar to those exposed to either visitor group, suggesting complementary effects of day and night pollination (Robertson et al. 2021). These findings highlight the need to understand how apple flowers interact with their pollinators across the full 24-hour cycle and whether floral traits have evolved to attract and reward nocturnal visitors as well as diurnal ones.

Successful pollination depends on the temporal coordination of floral rewards and pollinator activity. Many plant species regulate nectar secretion and scent emission through circadian rhythms that align with their pollinators’ foraging schedules (Edge et al. 2012, Fenske and Imaizumi 2016, Prieto-Benítez et al. 2016, Yan et al. 2016, Bloch et al. 2017, Fenske et al. 2018). Some nocturnally pollinated flowers are known to have compounds in their volatile bouquets that specifically attract night-active flower visitors (Carvalho et al. 2012, Cordeiro et al. 2017, Martínez-Martínez et al. 2021). If apple flowers emit volatiles and produce nectar during the night as well as the day, this would suggest an adaptive mechanism to attract pollinators from both temporal guilds. Abiotic factors such as temperature, cloudiness, humidity, and wind speed, may further modulate these dynamics by influencing both floral physiology and insect behavior (Holyoak et al. 1997, Vicens and Bosch 2000, Silva et al. 2004). Understanding how these biotic and abiotic processes interact is key to characterizing the temporal structure of apple pollination. Furthermore, the optimal environmental conditions for pollinator foraging, namely calm, warm, and cloudless weather, often coincide with those preferred for pesticide application (Karbassioon and Stanley 2023). This overlap underscores the need to define pesticide application windows that pose the least risk to beneficial insects, including the nocturnal pollinator guild.

The present study examines diel patterns in interactions between apple flowers and their potential pollinators. Specifically, we ask whether apple flowers recruit pollinators throughout the 24-h cycle. We quantify temporal variation in floral volatile composition and nectar production across multiple time points throughout the day and night and complement these measurements with video recordings of insect visits to flowers. As nocturnal pollination in apple remains understudied, we also use visual surveys and light trap collections to assess the presence of potential nocturnal pollinators during the nighttime periods, conducted concurrently with sampling flowers for volatiles and nectar. We also test whether environmental conditions influence both floral attractants and insect behavior. This integrative approach provides a comprehensive view of apple pollination ecology and offers practical insights for optimizing fruit production while conserving pollinator communities active across the entire diel cycle.

## 2. Materials and Methods

### 2.1. Study site

The study was conducted in an apple orchard at the Milo J. Shult Agricultural Research and Extension Center, University of Arkansas, Fayetteville (36°06′05″ N, 94°10′00″ W), during three bloom seasons: April 8–23, 2021, April 12–30, 2022, and March 31 – April 13, 2026. The 0.5-hectare orchard comprised ten rows with approximately 222 apple trees (*Malus domestica* Borkh.) of different varieties, including cultivars ‘Enterprise’ and ‘GoldRush’ as well as experimental breeding lines AR 124 and AR 127. The orchard was situated in a semi-rural environment, bordered by crop and fallow fields, and located about 100 m from a moderately busy road. Local light pollution at night was negligible, ensuring that the light traps used in this study were not affected by competing light sources. To ensure clarity regarding our temporal sampling structure, we explicitly distinguish between discrete sampling events and continuous monitoring intervals. Throughout this study, we define a time point as an instantaneous sampling event occurring at a specific hour (used for nectar collection, floral volatile collection, and visual transect surveys). Conversely, we define a time period as a continuous data collection interval spanning multiple hours (used for video recordings and light-trap sampling).

### 2.2. Weather records

Air temperature (°C), wind speed (m/s), and cloud cover (fair/cloudy) were recorded at each sampling time point and later verified using hourly weather data for the Drake Field airport (36°00′18″ N, 94°10′12″ W), available from the Integrated Surface Database (National Centers for Environmental Information (https://www.ncei.noaa.gov/products/land-based-station/integrated-surface-database). For continuous video recording, we used hourly air temperature (°C), wind speed (m/s), and cloud cover (%) directly from the weather database. No fieldwork was conducted in the rain.

### 2.3. Flower nectar collection and measurement

Nectar was collected during the 2021 season. To mimic the conditions experienced by potential nocturnal and diurnal pollinators when encountering apple flowers and their nectar rewards, we collected nectar from newly opened, unpollinated flowers that were not bagged prior to sampling. Nectar was collected at five sampling time points, weather permitting: two during the day (11:00 and 16:00), two at night (21:00 and 2:00), and one at dawn (6:00). At each time point, five blooming apple trees were randomly selected, and nectar was collected from five randomly chosen flowers per tree using 1-μL Drummond calibrated microcapillary tubes. Flowers were standardized across observers and sampling time points; they were freshly opened and did not show any visible signs of having been pollinated. Nectar was collected by inserting a microcapillary tube into the base of each flower for 1 min. Nectar volume was measured under a dissecting microscope against a reference scale prepared by dividing the length of a microcapillary tube into ten equal segments.

### 2.4. Flower volatiles collection and analysis

Apple flowers were collected for volatile extraction in 2021 and 2022 at five time points: two during the day (11:00 and 16:00), two at night (21:00 and 2:00), and one at dawn (6:00). At each sampling time point, two newly opened flowers from the same tree were cut off the stalk with clean dissecting scissors and placed in an amber-glass scintillation vial. Three randomly selected trees were sampled per time point, with each sample consisting of two flowers. Flowers from each tree were crushed together using a clean, sterilized glass rod, and the vial was immediately capped and sealed with parafilm. We sampled entire flowers as proxies for stored and emitted pools of metabolites because the apple orchard in situ experienced highly variable conditions that made headspace volatile collection challenging and interpretation difficult. We acknowledge that this sampling technique does not capture floral headspace, may capture wound-released metabolites, and relative proportions of compounds from crushed flowers may not perfectly match those released naturally by flowers (Jakobsen and Christensen 2002, Ibanez et al. 2010, Li et al. 2023). However, standard headspace collection techniques can also fail to capture all volatiles detected and used by insects (Nieberding et al. 2008, Visser et al. 2018). Using a crushed flower technique should increase the likelihood of capturing all the possible insect-detected compounds in the flower at the time of sampling. Samples were transported on ice to the laboratory and stored at 4 °C until analysis. Volatile analysis was conducted on a Shimadzu QP2020 gas chromatograph–mass spectrometer (GC–MS) equipped with a 30 m Restek Rtx-VMS capillary column (0.32 mm inner diameter, 1.8 µm film thickness). The oven temperature was ramped from 35 °C to 120 °C at a heating rate of 10°C/min and held for 10 minutes.

### 2.5. Flower visitation by insects

We documented insect visits to apple flowers across the 24-hour cycle using infrared video recordings in 2026. We also focused specifically on nighttime insect visitation, using a combination of infrared video recordings (2021), moth surveys along transects (2021 and 2022), and light-trap collections (2021 and 2022).

For video recording, one to four (usually two) full-spectrum night vision cameras (GhostStop, Lynchburg, VA, United States) were simultaneously deployed in the orchard. Cameras were mounted 1–2 m above ground and aimed at flower clusters containing multiple fresh or newly opened flowers. From dusk to dawn, the flower clusters were illuminated by battery-operated infrared lights. Videos were processed with the motion-detection software DeepMeerkat (Weinstein 2018) and then reviewed manually. All arthropods that contacted apple flowers were identified to a large taxonomic group (honeybees, bumblebees, other bees, wasps, brachyceran flies, nematoceran flies, moths, and miscellaneous), counted, binned into 2-h intervals, and later consolidated into five time periods centered on the timing of nectar and volatile sampling (0:00–4:00, 4:00–8:00, 8:00–14:00, 14:00–20:00, and 20:00–0:00).

Each transect survey in the orchard lasted 5 min and was carried out at 4 sampling time points: 18:00, 22:00, 2:00, and 6:00, during which all moths encountered along the transect were counted. For light trap collections, three pairs of quarter-acre insect traps (covering an area of approximately 1000 m^2^; DynaTrap, Lancaster, PA, United States) were installed at different locations within the orchard. Each pair was powered by a rechargeable battery to ensure continuous operation during sampling. Trap collections were conducted in three time periods: 18:00–22:00, 22:00–02:00, and 02:00–06:00. Only one pair of traps was active during each time period, such that a different orchard location was sampled every four hours. Insects captured during each time period were transferred to a zip-lock bag, stored at –20 °C, and later sorted and identified to family level.

### 2.6. Statistical analyses

All data processing, statistical analysis, and visualization were done in R v. 4.6.1 (R Core Team 2026) using RStudio v. 2026.08.0 (Posit Team 2026). Univariate response variables, such as nectar volume and total insect counts, were analyzed using generalized linear mixed-effects models (GLMMs). For nectar volume data, we fitted GLMMs by maximum likelihood in the package nlme, which is tailored to accommodate for heterogeneous variances and unequal sample sizes (Pinheiro and Bates 2000, Pinheiro et al. 2026). For insect count data, which contained a high proportion of zeros, we fitted Poisson GLMMs using glmmTMB (Brooks et al. 2017). Overall effects of model terms were assessed using likelihood-ratio tests (LRT), and differences among sampling time points or time periods were tested with Tukey-adjusted pairwise comparisons in the package emmeans (Lenth and Piaskowski 2026). Multivariate (compositional) responses, such as changes in floral volatile bouquets or insect community, were analyzed using permutational multivariate analysis of variance in the package vegan (Oksanen et al. 2026). Plots were generated with the package ggplot2 (Wickham 2016).

#### 2.6.1. Nectar

The effects of time of day (sampling time point), temperature, wind speed, and cloud cover on flower nectar volume were assessed using a linear mixed-effects model. Replicate apple trees nested within calendar date were included as random effects. Nectar data were log-transformed [ln(1+x)] to normalize the residuals, and the variance function structure allowed different variances for each time point, in order to account for heteroscedasticity. Model fit was evaluated using AICc scores, and the significance of effects was tested with likelihood-ratio chi-squared tests (LRT χ^2^). To visualize patterns of nectar volume across time of day and air temperature, mean values were calculated per tree.

#### 2.6.2. Volatiles

We reviewed the list of detected volatile and other metabolite compounds and removed likely contaminants, namely, silicon-containing compounds (dimethylsilanediol and various siloxanes), methylene chloride, and ethanol. Siloxanes and their derivative silanediol are manmade, and methylene chloride, while occurring naturally, is widely used as a solvent. Although ethanol can occur naturally in apple flowers, its high abundance suggested contribution from residual ethanol used for tool sterilization. We then quantified differences in volatile composition among samples in three complementary ways.

First, for descriptive visualization, we used log-transformed total ion counts (TIC) for selected volatiles and metabolites that were consistently detected across sampling time points and years and/or previously reported from apple floral scent (Loughrin et al. 1990, Buchbauer et al. 1993, Bengtsson et al. 2001, Fan et al. 2019, Rachersberger et al. 2019, Henneberg et al. 2022). These selected compounds were used for visualization only and not for the multivariate analyses of the complete volatile repertoire.

Second, relative abundances were calculated for all compounds within each sample and variation in volatile composition was analyzed using the R package chemodiv (Petrén et al. 2023, 2024). Bray-Curtis dissimilarity was used to characterize differences in compound identity and relative abundance, whereas Generalized UniFrac dissimilarity incorporated both relative abundance and structural dissimilarity among compounds. Differences in volatile composition among samples were evaluated using sequential and marginal PERMANOVA with sampling time point, sampling date, temperature, wind speed, and cloud cover as predictors. Since we were primarily interested in the effect of time of day, the sequential model was specified as *D*(*Y*) ∼ time point + date + temperature + wind speed + cloud cover. Marginal tests of the same predictors were used as a robustness check. We also visualized variation among volatile samples collected at the five time points using principal coordinates analysis (PCoA).

As time of day significantly affected volatile composition in 2021, we conducted a follow-up constrained ordination to characterize this effect. We fitted a partial distance-based redundancy analysis (dbRDA) to the 2021 Generalized UniFrac distance matrix, with time of day as the constrained factor and calendar date, temperature, wind speed, and cloud cover included as conditioning variables. Square-root-transformed relative abundances were added to the partial dbRDA as supplementary compound scores. For each compound, we calculated the Euclidean magnitude of its scores across all constrained dbRDA axes and ranked compounds by this value. These scores were used to identify compounds most strongly associated with the time-of-day-constrained ordination.

Finally, we used a compound-free approach to compare floral volatile bouquets, applying the “Compounds Without Borders” (CWB) framework (Sprayberry 2020). This method characterizes volatiles based on structural features (e.g., total carbon count, chain length, functional groups, and cyclic carbons) and calculates angular distances between treatments in a 66-dimensional space. Larger angular values indicate greater chemical dissimilarity between treatments. For this analysis, we first averaged total ion counts for each compound within a sample and expressed them as percentages. We then calculated mean compound percentages per time point and used these values as inputs for the CWB analyses.

#### 2.6.3. Insect visitation from video recordings

The duration of usable footage varied among 2-h time periods, and so we calculated insect counts per hour of footage. Differences in insect community composition were quantified using Bray-Curtis dissimilarity. The effects of time of day, calendar date, temperature, wind speed, and cloud cover were evaluated using sequential and marginal PERMANOVA. Overall community patterns were visualized using vertical bar charts of raw insect counts over time and nonmetric multidimensional scaling (NMDS) of insect counts per hour.

#### 2.6.4. Other nocturnal insects in the orchard

We tested the effect of time of night on total insect counts in transect surveys and light traps using a one-way Kruskal-Wallis ANOVA, followed by a pairwise post-hoc Wilcoxon test with a Bonferroni correction, separately for each year and sampling method. As both sampling methods yielded consistent results throughout the nighttime sampling period, we pooled these datasets together and fitted a generalized linear mixed-effects model with insect count as the response variable and year, time of night, temperature, wind speed, and cloud cover as predictor variables. To account for systematic baseline differences across sampling method, the latter was treated as a random intercept.

## 3. Results

### 3.1. Weather conditions

Spring weather differed markedly between the sampling years, both prior to and during the sampling season, with corresponding shifts in apple phenology. In 2021 and 2026, March was relatively warm (monthly means 11.2 and 15.5 °C, respectively), and apple trees began blooming in late March. In contrast, March 2022 was cooler (mean 9.1 °C), and bloom onset was delayed. As a result, sampling in 2021 (April 8–23) coincided with the peak and latter, declining phase of bloom, whereas sampling in 2022 (April 12–30) and 2026 (March 31 – April 13) encompassed both the onset and peak of bloom.

Weather conditions also differed during the sampling periods themselves. The 2021 sampling period was the coolest, with average daily mean, minimum, and maximum temperatures of 11.4, 5.5, and 17.2 °C, respectively, whereas temperatures in 2022 (15.5, 9.8, and 20.9 °C) and 2026 (18.2, 11.9, and 24.6), were slightly and considerably above the multi-year norm, respectively. Precipitation also contrasted sharply: April 2021 and 2026 were relatively dry, with only 43.9 and 73.2 mm of rainfall during the sampling period (about 40% and 70% of the long-term monthly average), whereas April 2022 was exceptionally wet, with 159.5 mm of rainfall (150% of the average). Wind speeds were lower in 2021 (3.9 m/s) and 2026 (4.3 m/s) than in 2022 (5.4 m/s). Approximately equal numbers of samples were taken (or surveys made) across cloudy and fair weather.

### 3.2. Nectar is consistently present in apple flowers throughout the diel cycle

In total, nectar was collected from 156 trees and 779 flowers (due to weather, we were unable to sample a complete set of 5 flowers from 5 trees on two occasions). Nectar volume in apple flowers varied considerably between individual samples but remained consistent across the diel cycle. Notably, at every time point except 16:00, some trees produced exceptionally high nectar volumes (Fig. 1). On average, nectar volume remained at 0.04–0.05 μL per flower at all the five sampling time points. A linear mixed-effects model showed no significant support for the effects of time of day (LRT χ^2^ = 0.1, *P* = 0.8), cloud cover (LRT χ^2^ = 1.0, *P* = 0.3) or wind speed (LRT χ^2^ = 2.9, *P* = 0.09), but a significant effect of air temperature (LRT χ^2^ = 7.6, *P* = 0.006), with higher nectar volumes observed under cooler conditions, particularly when temperature was 10 °C or lower (Supplementary Fig. S1).

**Fig. 1.**
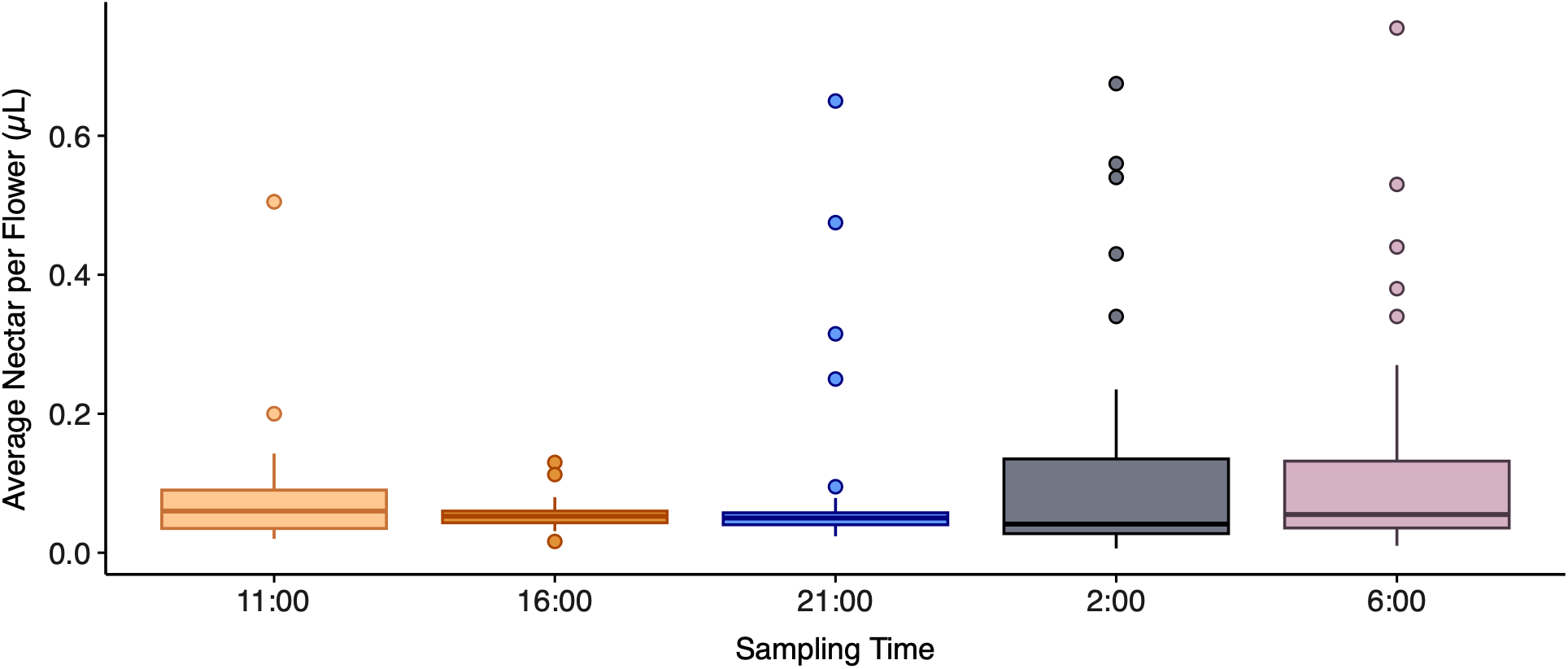
Variation in floral nectar volume in apple flowers: nectar production remained stable across five sampling times during the day and night. Horizontal lines denote medians, boxes show the interquartile range, and whiskers extend to 1.5 × the interquartile range. Results of the mixed-effects linear model are reported in the text.

### 3.3. Floral volatiles are present throughout the diel cycle but vary with time of day

In total, 89 floral volatile samples were collected in 2021 and 98 in 2022 (due to weather, we were unable to sample from all the three trees per time point on two occasions). The 2021 samples comprised 130 volatile and other metabolite compounds (Supplementary Table S1) as well as ethanol, methylene chloride, and three siloxanes, which we excluded. The 2022 samples comprised 126 compounds (Supplementary Table S2) plus ethanol, dimethylsilanediol, and ten siloxanes, which we excluded. To illustrate the major constituents of the apple floral volatile bouquet, we focused on 19 compounds that were frequently detected across samples and/or are known components of apple floral scent. These compounds consisted primarily of aldehydes, particularly medium-chain aliphatic aldehydes, followed by alkanes and alcohols (Fig. 2A).

**Fig. 2.**
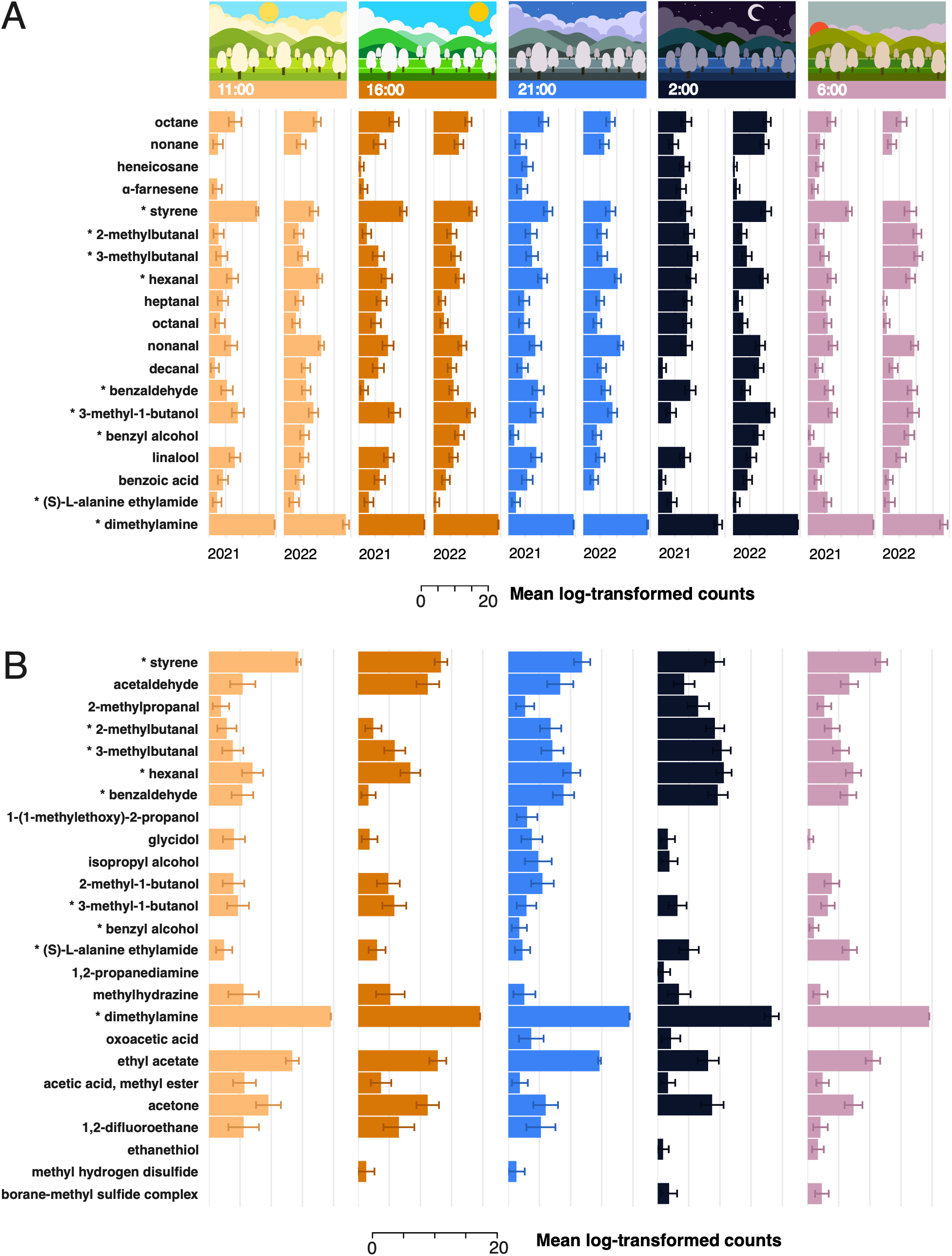
Diel variation in floral volatile composition of apple flowers, shown as mean (± SE) log-transformed ion counts of selected compounds. (A) Predefined set of floral volatiles of interest that were consistently detected across sampling times and years and/or had been previously reported as components of apple floral scent. (B) Compounds most strongly associated with the sampling-time-constrained ordination, ranked by the Euclidean magnitude of their supplementary scores across all constrained dbRDA axes after accounting for calendar date, temperature, wind speed, and cloud cover. A complete ranking of compounds is provided in Supplementary Table S3. Asterisks denote compounds present in both (A) and (B).

We then used the entire set of detected compounds (except organosilicons, methylene chloride, and ethanol) and fitted full PERMANOVA models including the effects of sampling year, time of day, temperature, wind speed, and cloud cover. These analyses revealed a strong effect of year on both volatile composition and chemical dissimilarity, as well as significant differences in multivariate dispersion (Supplementary Fig. S2). We therefore analyzed the 2021 and 2022 volatile datasets separately.

Volatile profiles varied significantly among sampling time points (times of day) in 2021 but not in 2022 (Table 1). In 2021, Generalized UniFrac dissimilarities, which account for both relative compound abundance and chemical structure, showed a significant effect of time of day in both sequential and marginal PERMANOVA. In contrast, Bray-Curtis dissimilarities, which primarily reflect differences in relative abundance, showed a significant time-of-day effect only in the sequential test (Table 1). Dispersion differed significantly throughout the day in 2021 for Bray-Curtis dissimilarity (*F* = 2.8, *P* = 0.03), but not for Generalized UniFrac (*F* = 1.9, *P* = 0.1, df = 4, 82). Thus, the inconsistent 2021 Bray-Curtis pattern (significant effect of time of day in sequential but not marginal PERMANOVA) may partly reflect differences in volatile data scatter within each sampling time point, whereas Generalized UniFrac differences clearly reflect significant separation of time-of-day centroids. In 2022, dispersion did not differ significantly among times of day for either Bray-Curtis (*F* = 0.3, *P* = 0.9) or Generalized UniFrac (*F* = 0.2, *P* = 0.06, df = 4, 93). Despite the significant effect of time of day detected by PERMANOVA, samples showed substantial overlap in PCoA ordination space, with no clear separation among sampling time points (Fig. 3A-D), although the spread of samples visible in the PCoA ordination for Bray-Curtis (Fig. 3A) was consistent with significant differences in multivariate dispersion among the sampling times. Overall, calendar date explained the largest proportion of variation among the factors included in the model in both years, and its effect was highly significant in most analyses (Table 1). Among the weather variables, only temperature had a weak effect on both relative-abundance and chemical composition, and only in 2021 (Table 1). Floral volatiles and other metabolites contributing strongly to time-of-day differences in 2021 included medium-chain aliphatic alcohols and aldehydes, benzenoids, and nitrogen-containing compounds (Fig. 2B; Supplementary Table S3). Nine of these compounds were in our initial list of consistently and abundantly detected volatiles throughout the day and across years (Fig. 2A).

**Fig. 3.**
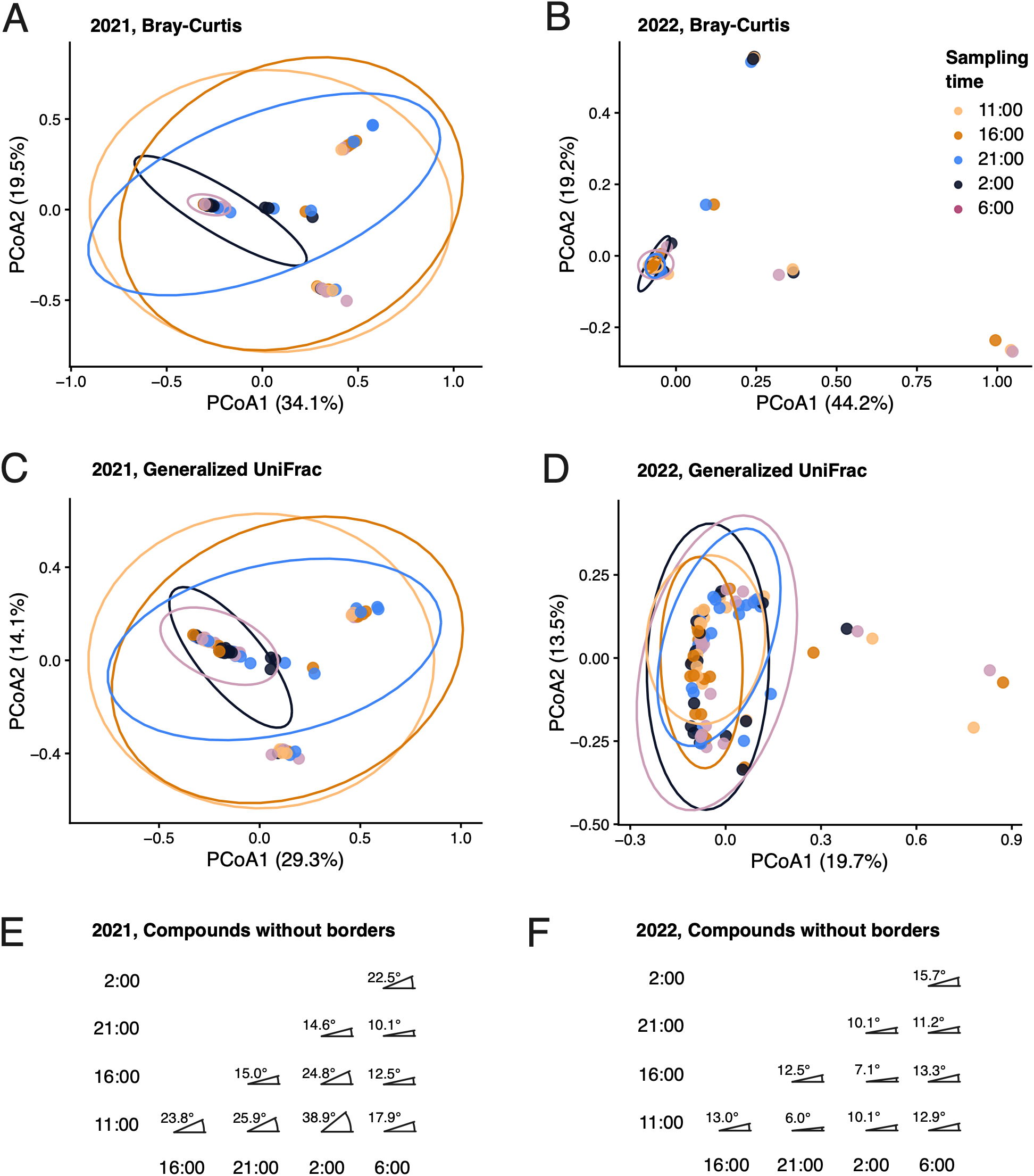
Multivariate patterns of variation in floral volatile bouquets across sampling time points in 2021 and 2022. Principal coordinates analysis (PCoA) ordinations are shown for (A) 2021 and (B) 2022 based on Bray-Curtis dissimilarity, and for (C) 2021 and (D) 2022 based on Generalized UniFrac dissimilarity incorporating molecular structural differences among compounds. Cailliez correction was applied to all PCoA distance matrices. Points represent individual samples and are colored according to sampling time point across the 24-h cycle; ellipses denote 95% confidence regions for each group. Compounds-Without-Borders (CWB) analysis shows angular differences in volatile bouquets among sampling times in (E) 2021 and (F) 2022.

**Table 1.**
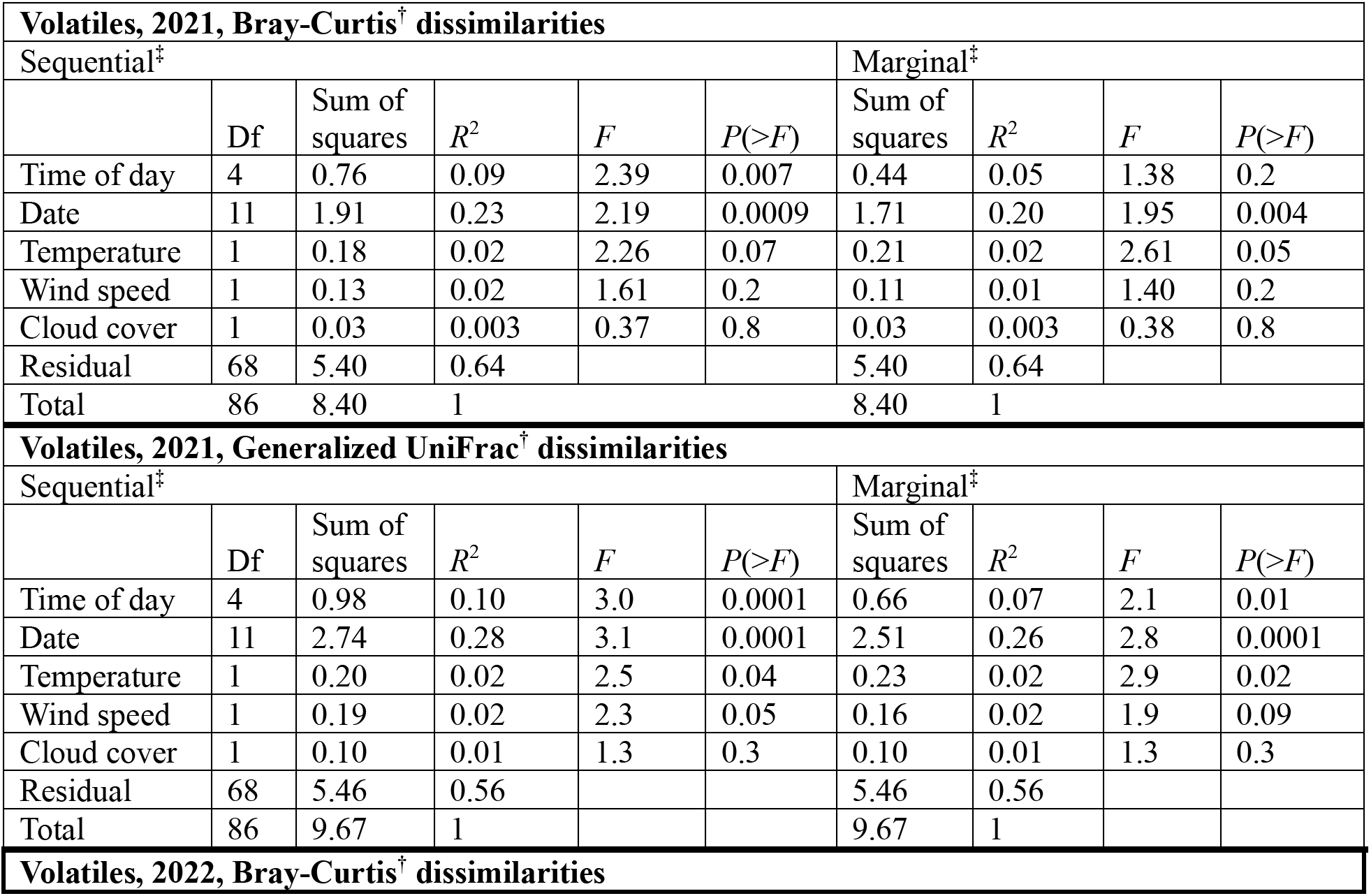
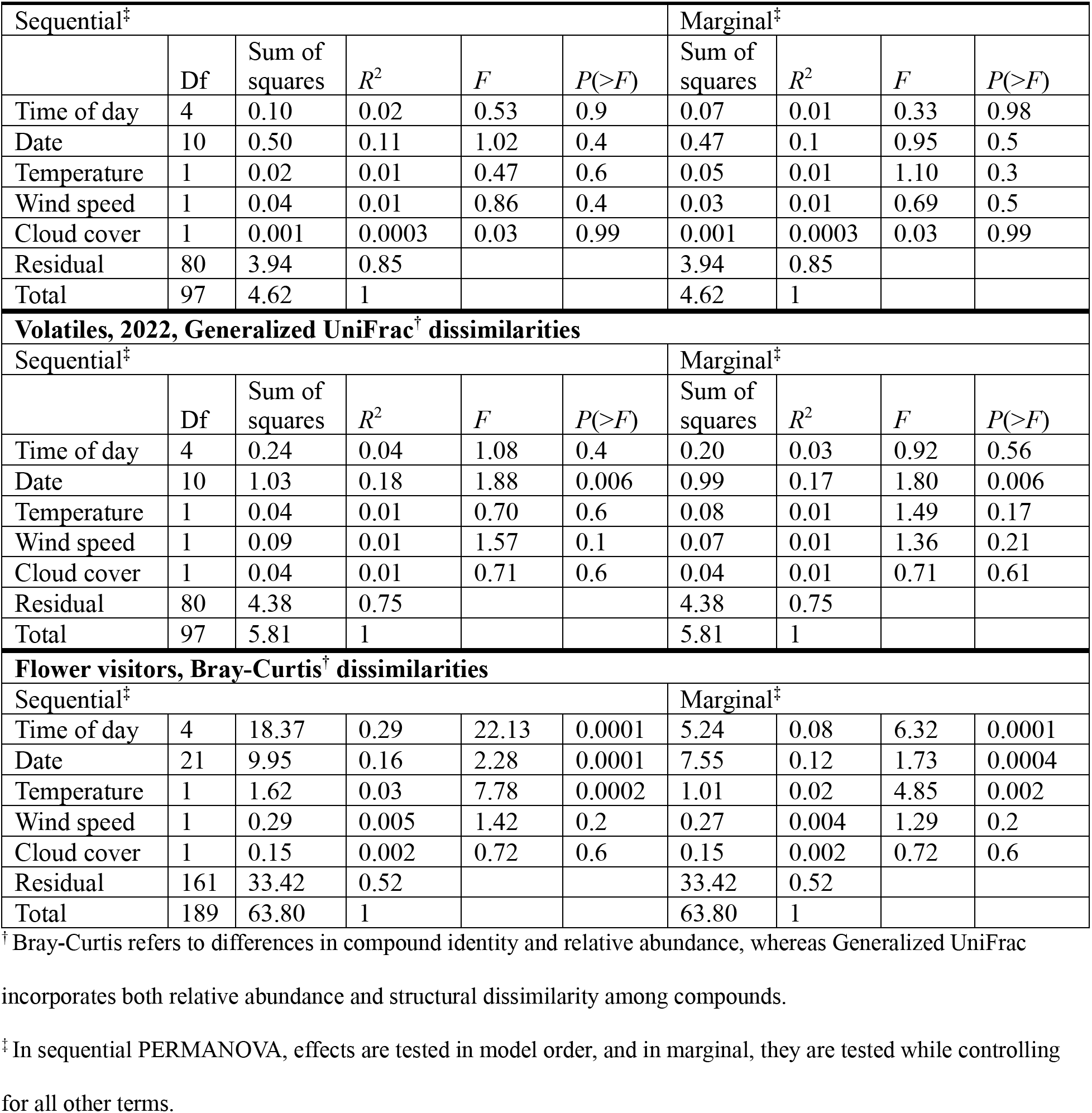
The effects of calendar date, time of day, temperature, wind speed, and cloud cover on apple floral volatile profiles and visitor community – results of permutational multivariate analyses of variance (PERMANOVA)

| **Volatiles, 2021, Bray-Curtis<sup>†</sup> dissimilarities** |  |  |  |  |  |  |  |  |  |
|---|---|---|---|---|---|---|---|---|---|
| Sequential<sup>‡</sup> |  |  |  |  |  | Marginal<sup>‡</sup> |  |  |  |
|  | Df | Sum of squares | *R*<sup>2</sup> | *F* | *P*(>*F*) | Sum of squares | *R*<sup>2</sup> | *F* | *P*(>*F*) |
| Time of day | 4 | 0.76 | 0.09 | 2.39 | 0.007 | 0.44 | 0.05 | 1.38 | 0.2 |
| Date | 11 | 1.91 | 0.23 | 2.19 | 0.0009 | 1.71 | 0.20 | 1.95 | 0.004 |
| Temperature | 1 | 0.18 | 0.02 | 2.26 | 0.07 | 0.21 | 0.02 | 2.61 | 0.05 |
| Wind speed | 1 | 0.13 | 0.02 | 1.61 | 0.2 | 0.11 | 0.01 | 1.40 | 0.2 |
| Cloud cover | 1 | 0.03 | 0.003 | 0.37 | 0.8 | 0.03 | 0.003 | 0.38 | 0.8 |
| Residual | 68 | 5.40 | 0.64 |  |  | 5.40 | 0.64 |  |  |
| Total | 86 | 8.40 | 1 |  |  | 8.40 | 1 |  |  |
| **Volatiles, 2021, Generalized UniFrac<sup>†</sup> dissimilarities** |  |  |  |  |  |  |  |  |  |
| Sequential<sup>‡</sup> |  |  |  |  |  | Marginal<sup>‡</sup> |  |  |  |
|  | Df | Sum of squares | *R*<sup>2</sup> | *F* | *P*(>*F*) | Sum of squares | *R*<sup>2</sup> | *F* | *P*(>*F*) |
| Time of day | 4 | 0.98 | 0.10 | 3.0 | 0.0001 | 0.66 | 0.07 | 2.1 | 0.01 |
| Date | 11 | 2.74 | 0.28 | 3.1 | 0.0001 | 2.51 | 0.26 | 2.8 | 0.0001 |
| Temperature | 1 | 0.20 | 0.02 | 2.5 | 0.04 | 0.23 | 0.02 | 2.9 | 0.02 |
| Wind speed | 1 | 0.19 | 0.02 | 2.3 | 0.05 | 0.16 | 0.02 | 1.9 | 0.09 |
| Cloud cover | 1 | 0.10 | 0.01 | 1.3 | 0.3 | 0.10 | 0.01 | 1.3 | 0.3 |
| Residual | 68 | 5.46 | 0.56 |  |  | 5.46 | 0.56 |  |  |
| Total | 86 | 9.67 | 1 |  |  | 9.67 | 1 |  |  |
| **Volatiles, 2022, Bray-Curtis<sup>†</sup> dissimilarities** |  |  |  |  |  |  |  |  |  |
| Sequential<sup>‡</sup> |  |  |  |  |  | Marginal<sup>‡</sup> |  |  |  |
|  | Df | Sum of squares | *R*<sup>2</sup> | *F* | *P*(>*F*) | Sum of squares | *R*<sup>2</sup> | *F* | *P*(>*F*) |
| Time of day | 4 | 0.10 | 0.02 | 0.53 | 0.9 | 0.07 | 0.01 | 0.33 | 0.98 |
| Date | 10 | 0.50 | 0.11 | 1.02 | 0.4 | 0.47 | 0.1 | 0.95 | 0.5 |
| Temperature | 1 | 0.02 | 0.01 | 0.47 | 0.6 | 0.05 | 0.01 | 1.10 | 0.3 |
| Wind speed | 1 | 0.04 | 0.01 | 0.86 | 0.4 | 0.03 | 0.01 | 0.69 | 0.5 |
| Cloud cover | 1 | 0.001 | 0.0003 | 0.03 | 0.99 | 0.001 | 0.0003 | 0.03 | 0.99 |
| Residual | 80 | 3.94 | 0.85 |  |  | 3.94 | 0.85 |  |  |
| Total | 97 | 4.62 | 1 |  |  | 4.62 | 1 |  |  |
| **Volatiles, 2022, Generalized UniFrac<sup>†</sup> dissimilarities** |  |  |  |  |  |  |  |  |  |
| Sequential<sup>‡</sup> |  |  |  |  |  | Marginal<sup>‡</sup> |  |  |  |
|  | Df | Sum of squares | *R*<sup>2</sup> | *F* | *P*(>*F*) | Sum of squares | *R*<sup>2</sup> | *F* | *P*(>*F*) |
| Time of day | 4 | 0.24 | 0.04 | 1.08 | 0.4 | 0.20 | 0.03 | 0.92 | 0.56 |
| Date | 10 | 1.03 | 0.18 | 1.88 | 0.006 | 0.99 | 0.17 | 1.80 | 0.006 |
| Temperature | 1 | 0.04 | 0.01 | 0.70 | 0.6 | 0.08 | 0.01 | 1.49 | 0.17 |
| Wind speed | 1 | 0.09 | 0.01 | 1.57 | 0.1 | 0.07 | 0.01 | 1.36 | 0.21 |
| Cloud cover | 1 | 0.04 | 0.01 | 0.71 | 0.6 | 0.04 | 0.01 | 0.71 | 0.61 |
| Residual | 80 | 4.38 | 0.75 |  |  | 4.38 | 0.75 |  |  |
| Total | 97 | 5.81 | 1 |  |  | 5.81 | 1 |  |  |
| **Flower visitors, Bray-Curtis<sup>†</sup> dissimilarities** |  |  |  |  |  |  |  |  |  |
| Sequential<sup>‡</sup> |  |  |  |  |  | Marginal<sup>‡</sup> |  |  |  |
| Time of day | 4 | 18.37 | 0.29 | 22.13 | 0.0001 | 5.24 | 0.08 | 6.32 | 0.0001 |
| Date | 21 | 9.95 | 0.16 | 2.28 | 0.0001 | 7.55 | 0.12 | 1.73 | 0.0004 |
| Temperature | 1 | 1.62 | 0.03 | 7.78 | 0.0002 | 1.01 | 0.02 | 4.85 | 0.002 |
| Wind speed | 1 | 0.29 | 0.005 | 1.42 | 0.2 | 0.27 | 0.004 | 1.29 | 0.2 |
| Cloud cover | 1 | 0.15 | 0.002 | 0.72 | 0.6 | 0.15 | 0.002 | 0.72 | 0.6 |
| Residual | 161 | 33.42 | 0.52 |  |  | 33.42 | 0.52 |  |  |
| Total | 189 | 63.80 | 1 |  |  | 63.80 | 1 |  |  |

<sup>†</sup> Bray-Curtis refers to differences in compound identity and relative abundance, whereas Generalized UniFrac incorporates both relative abundance and structural dissimilarity among compounds.
<sup>‡</sup> In sequential PERMANOVA, effects are tested in model order, and in marginal, they are tested while controlling for all other terms.

Compounds Without Borders (CWB), which quantifies angular differences between treatments in a 66-dimensional space based on identified compound characteristics (Sprayberry 2020), revealed stronger contrasts between volatiles sampled at different times in 2021 than in 2022. In 2021, the largest angular separations occurred between volatiles released at night (21:00 and 2:00) and those released during the day (11:00 and 16:00; Fig. 3E). By contrast, in 2022, angular values were mostly lower than those observed in 2021 across all time-point comparisons (Fig. 3F).

### 3.4. Different groups of insects visit apple flowers around the clock

We processed 431 h of usable video footage collected across all cameras and both years, representing 235 two-hour sampling periods. In total, we recorded 2,711 arthropod visits to apple flowers: 889 honeybees, 16 bumblebees, 624 other bees, 54 wasps, 647 brachyceran flies, 81 nematoceran flies (e.g., a cranefly in Supplementary Video 1), 165 moths (e.g., a hawkmoth in Supplementary Video 2), and 235 miscellaneous visitors, including beetles, lacewings, ants, spiders, true bugs, butterflies, and unidentified insects, many of which were likely small flies (see Supplementary Fig. S3 for additional photos and Fig. S4 for video screenshots of some flower visitors). Hymenoptera and Brachycera visited flowers only between 06:00 and 20:00, whereas Nematocera and moths were primarily active between 18:00 and 06:00 and were rarely observed during the day (Fig. 4A).

**Fig. 4.**
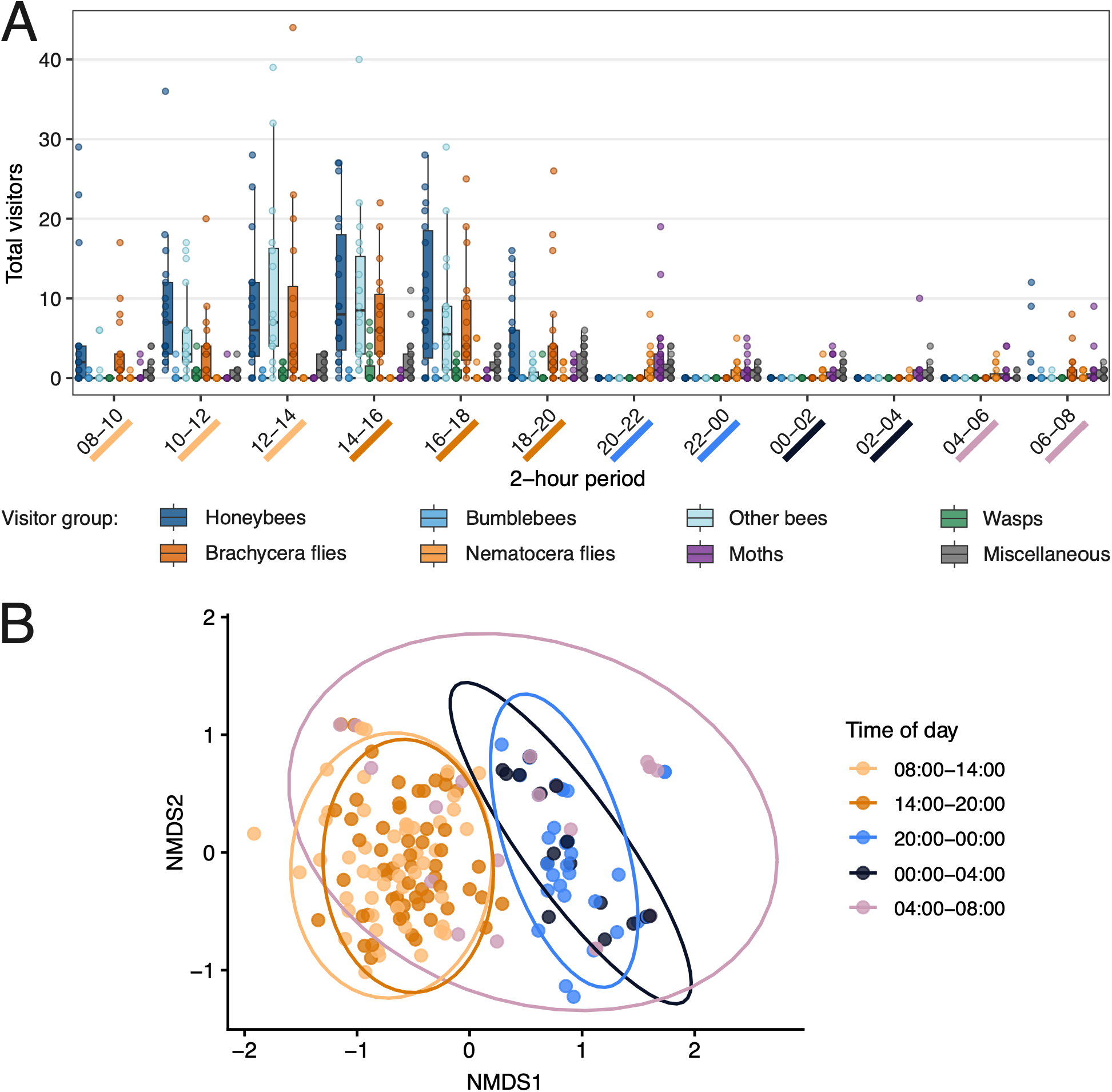
Diel variation in apple flower-visiting insect abundance and community composition, 2021 and 2026 video recordings combined. (A) Total raw insect counts (unstandardized by the actual duration of usable footage) per 2-h video-recording period, broken down by visitor taxonomic group. Each data point represents a unique combination of visitor group, flower cluster, calendar date, and 2-h recording period. Horizontal lines indicate medians, boxes show the interquartile range (IQR), and whiskers extend to 1.5 × IQR. (B) Non-metric multidimensional scaling (NMDS) ordination of visitor community composition across five diel sampling periods. For this analysis, consecutive 2-h video-recording periods were consolidated into broader time intervals centered on the five sampling times used for nectar and floral volatile collections. Points are colored by sampling period, with proximity among points indicating greater similarity in visitor community composition. Ellipses denote 95% confidence regions for each sampling-period group.

As some periods contained less than 2 h of footage, visitation rates were standardized as the number of visitors per hour. We first fitted full PERMANOVA models including the effects of sampling year, time of day, temperature, wind speed, and cloud cover on the taxonomic composition of insect visitors. The effect of year was nonsignificant, so the 2021 and 2026 data were pooled for subsequent analyses (separate bar plots of raw insect counts for each year are shown in Supplementary Fig. S5). Flower visitor community composition was significantly influenced by time of day, calendar date, and temperature (Table 1). NMDS ordination showed a clear temporal pattern: the two daytime periods clustered closely together, as did the two nighttime periods, while early morning (04:00–08:00) was intermediate between them (Fig. 4B).

### 3.5. Nighttime insect activity in the orchard is weather-dependent and declines by early morning

We surveyed a total of 50 transects in the apple orchard: 24 in 2021 and 26 in 2022. Along these transects, we recorded 104 moths: 29 individuals over four sampling time points in 2021 and 75 over three sampling time points in 2022. We collected 42 insect samples from light traps: 18 in 2021 and 24 in 2022, each sample representing a total catch after a four-hour operation of two traps. These yielded a total of 353 individuals, 65 of which were in 2021 and 288 in 2022. Craneflies dominated the catch (190 individuals: 186 Tipulidae and 4 Limoniidae), nearly all from 2022 (182 individuals). We also collected 36 moths (18 in each year), 125 insects from other taxa (39 in 2021 and 86 in 2022), and 4 spiders (all in 2022). The diversity and abundance of taxa caught in light traps are shown in Supplementary Fig. S6 and S7. Insect abundance and diversity differed sharply between years. We therefore fitted separate GLMMs for each sampling method and year. Across both years and sampling methods, insect abundance declined over the course of the night, with few or no insects recorded after 2:00, and increased significantly with rising temperature (Supplementary Table S4; Supplementary Fig. S8 and S9). Wind speed was positively associated with insect abundance in 2022 for both sampling methods, whereas cloud cover had a significant effect only in 2022 light-trap samples (Supplementary Table S4). The latter pattern was driven largely by a single cloudy night with particularly high cranefly abundance.

## 4. Discussion

Using three complementary lines of evidence (nectar production, floral volatiles, and insect visitation patterns), this study demonstrates that apple trees in Northwest Arkansas remain attractive to potential pollinators across the 24-hour diel cycle. First, nectar volume remained relatively constant over a 24-hour period, indicating that apple flowers invest similarly in both diurnal and nocturnal pollination. Second, crushed flowers contained comparable amounts of volatiles and other metabolites throughout the day-night cycle, while volatile composition exhibited diel variation, with the most pronounced changes occurring during the transition from darkness to dawn to daylight – precisely when insect visitor abundance was lowest. Third, the Hymenoptera-dominated daytime flower visitor community was replaced by moths at night, whereas Diptera were present on flowers around the clock, with Brachycera during the day and Nematocera at night. Weather effects were generally small and rarely significant, but nectar volume decreased and insect abundance increased with rising temperature.

Our findings show that nocturnal visitation by insects is unlikely to be incidental; instead, apple flowers appear to allocate resources – including floral volatiles and accessible nectar – toward attracting and rewarding visitors at night. The primary daytime pollinators of apple are bees – both managed honeybees and diverse wild species (Stern et al. 2001, Ramírez and Davenport 2013, Földesi et al. 2016, Park et al. 2016, Quinet et al. 2016); this study). But, accumulating evidence indicates that nocturnal insects also contribute meaningfully to pollen transfer and seed set (Robertson et al. 2021, Chatterjee and Singh 2025), and our data suggest this diel recruitment of possible pollinators is facilitated by apple flower attractants and rewards. As efforts to include pollinator protection into apple integrated pest management continue (Phan et al. 2025), our results suggest that these guidelines may need to expand to explicitly account for the nocturnal pollinator community.

We observed diel shifts in crushed apple flower volatiles to be driven largely by quantitative changes in aldehydes, alkanes, and benzenoids, a pattern consistent with differential attraction of day-versus night-active flower visitors. Interannual shifts in volatiles from crushed apple floral tissues also mostly involve aldehydes, alkanes, and benzenoids, and may reflect the substantial weather differences between 2021 and 2022. Similar rainfall- and drought-driven changes in floral volatile emissions have been documented across diverse systems (Campbell et al. 2019, Farré-Armengol et al. 2020, Rering et al. 2020, Powers et al. 2025). Given that diel, interannual, and weather-driven shifts in volatiles involve changes in similar types of chemical compounds, it would be interesting to see if the observed interannual and weather-driven shifts in volatiles are also associated with interannual or weather-driven shifts in floral visitor assemblages.

The volatile bouquet we recovered from crushed apple blossoms strongly suggests the flowers are attracting Diptera and moths, not just bees. Floral volatiles typically comprise varying proportions of fatty acid derivatives, terpenoids, and benzenoids, with one group usually dominating the profile (Dobson 2006). In our apple samples, fatty acid derivatives are most abundant, followed by benzenoids (styrene, benzoic acid, benzaldehyde, and benzyl alcohol) and a smaller fraction of terpenoids (heneicosane, α-farnesene, and linalool). While the volatile composition of floral tissues, as analyzed here, does not necessarily reflect the identity or relative abundance of compounds released into the floral headspace (Jakobsen and Christensen 2002, Ibanez et al. 2010, Li et al. 2023), large quantities of fatty acid derivatives and benzenoids are characteristic of Diptera- and moth-visited flowers (Shaver et al. 1997, Dobson 2006, Farré-Armengol et al. 2020) and are prominent in scented Myrtaceae fruit crops (Cordeiro et al. 2019). Alkanes and nitrogenous compounds, which are particularly abundant in Diptera-pollinated flowers (Dobson 2006, Cardoso-Gustavson et al. 2017, Zhang et al. 2018, Arnold et al. 2019), are likewise represented here, including alanine ethylamide. Several of these apple floral volatiles also exhibit diel rhythmicity, with nighttime bouquets differing from those present during the day and at dawn. One nitrogen-containing compound, dimethylamine, is particularly abundant across all conditions. Blossom tissues of other pome-bearing Rosaceae, such as *Crataegus* and *Sorbus*, are likewise known to be rich in amines (Sintermann and Neftel 2015), and these flowers are frequently visited by flies (Fichtner and Wissemann 2021).

Several of the compounds we observed are known to elicit nocturnal insect responses, e.g., benzyl alcohol, α-farnesene, linalool, methylbutanols, and methylbutanals (Shaver et al. 1997, Jürgens et al. 2002, Chen et al. 2015). Codling moths, for example, show antennal sensitivity to α-farnesene, benzyl alcohol, linalool, and methyl salicylate released from various apple tissues (Bengtsson et al. 2001). In our 2021 dataset, three of these (α-farnesene, benzyl alcohol, and methyl salicylate) are most abundant at night, whereas linalool is detected more uniformly across the diel cycle in both years. The enantiomeric composition of linalool, which can mediate species-specific communication and reduce herbivore attraction (Raguso 2016), also raises the possibility that day–night shifts in linalool enantiomers could serve as a private communication channel for pollinator interactions – an intriguing avenue for future work. Many of these moth-attractive compounds also occur at high levels in night-blooming *Silene* species (Jürgens et al. 2002), underscoring their broader role in nocturnal pollination systems. In crabapple (*M. ioensis*) flowers, emissions peak in the afternoon and are enriched in alcohols and ketones, whereas nighttime samples contain more methyl hexadecanoate and dawn/dusk samples are elevated in α-pinene (Fan et al. 2019). Our findings reveal a comparable temporal organization, which is more fine-scale than broad day-night differences. Interestingly, none of the compounds most strongly associated with the sampling-time-constrained ordination were unique to daytime samples, whereas six of these compounds occurred only at night or also at dawn.

Volatile compound identities should be interpreted cautiously because assignments were based on the best GC-MS library matches and were not confirmed with analytical methods. That said, many detected compounds in our samples match those reported previously, but their relative proportions differ, potentially reflecting methodological differences in volatile recovery. Our samples most closely resemble solvent-extracted bouquets (Omata et al. 1990, Buchbauer et al. 1993). By contrast, apple headspace studies often detect higher proportions of benzenoids – particularly benzyl alcohol and/or benzaldehyde (Loughrin et al. 1990, Buchbauer et al. 1993, Piskorski and Dorn 2010) – although exceptions exist, such as the predominance of both benzyl alcohol and α-farnesene (Bengtsson et al. 2001) or α-farnesene and linalool (Fraternale et al. 2014). It would be interesting to see if head space collections taken at night also recover high proportions of fatty acid derivatives in comparison to daytime samplings.

Nocturnal pollination is increasingly recognized as an important component of plant reproductive ecology, but the identity and effectiveness of nocturnal visitors remain understudied. A recent global meta-analysis showed that in ∼90% of angiosperms tested, diurnal and nocturnal pollination were similarly effective, and flowers exposed continuously for 24 h generally had the highest pollination success (Kendall and Nicholson 2025). Apple fits this broader pattern: the pollinator exclusion experiment conducted at the same facility as the present study (Robertson et al. 2021) was part of that meta-analysis, and its results – significant nocturnal contributions to seed set – align well with the global synthesis. However, the comparable seed set among flowers exposed only during the day, only at night, or across the full diel cycle suggests that at least one group of potentially effective apple pollinators might operate throughout the 24-hour period. Dipterans, which are abundant and often inconspicuous, represent a plausible candidate for such around-the-clock activity. This interpretation is consistent with our video observations: brachyceran flies were abundant daytime visitors, whereas nematoceran flies were common at night, and many of the small insects classified as “Miscellaneous” were likely additional Diptera that could not be identified reliably from the footage; still smaller visitors may have gone undetected altogether because the video resolution was insufficient for consistent detection and identification of very small insects.

Indeed, dipterans are common nocturnal visitors in other fruit crops: in blueberries, nocturnal insects captured near flowers were dominated by Diptera, followed by moths (Cutler et al. 2012). Although their pollination efficiency varies widely, many dipteran families – including several found in our light traps (Tipulidae, Chironomidae, Mycetophilidae, Culicidae, Psychodidae, and Anthomyiidae) – are well documented as visitors to apple blossoms (Wilson 1929, Kendall and Solomon 1973, Pansarin and Pansarin 2017) and/or confirmed pollinators in other systems (Larson et al. 2001), attracted by floral odors and nectar rewards (Woodcock et al. 2014). Hoverflies (Syrphidae) are also prominent diurnal apple pollinators (Wilson 1929, Kendall and Solomon 1973, Burns and Stanley 2022, McCravy et al. 2024, Mupepele et al. 2025). Moths likewise constitute an important group of nocturnal flower visitors and potential pollinators. Although moth visitation to apple blossoms has been less frequently documented than for Diptera (but see Robertson *et al*., 2021), recent work has recorded several moth families – including Noctuidae, Erebidae, and Geometridae – on apple flowers and identified them as particularly effective nocturnal pollen vectors (Chatterjee and Singh 2025). Moths caught in our light traps were primarily Plutellidae, Noctuidae, Crambidae, and Gelechiidae, while video recordings primarily documented flower visits by Sphingidae and Noctuidae; as with Diptera, we likely underestimated the visits of smaller moths to apple flowers. Insect activity was lowest on cooler nights (below 10°C) and generally declined as the night progressed, suggesting that – when pesticide applications during bloom are unavoidable – the ecologically least damaging window may be on cooler nights after 2:00, provided this timing is practical for growers. Any such recommendation, however, must be weighed against potential reductions in pesticide efficacy and the operational constraints of commercial orchards.

Nectar was consistently present in apple flowers over the diel cycle. While the quantity of nectar we observed per flower may seem low, it falls well within the wide range previously reported for apple flowers (Southwick et al. 1981, Abrol 1990, Quinet et al. 2016, Smessaert et al. 2019). Apple nectar volume can also vary greatly from year to year (Schneider et al. 2002) and with floral age (Farkas and Zajácz 2007). In the present study, we intentionally did not bag flowers prior to nectar sampling, as our aim was to evaluate nighttime nectar availability under conditions that allowed nectar removal by daytime visitors. However, even bagged flowers – at least in pears, whose nectaries are more exposed than those of apples – show considerable fluctuations in nectar volume, likely due to dilution by morning dew and subsequent evaporation or resorption during the day (Sponsler et al. 2024). Consequently, the negative relationship we observed between nectar volume in freshly opened apple flowers and air temperature may reflect both abiotic (direct effect of temperature) and biotic influences (increased insect activity). Even with these limitations in mind, our results indicate that apple flowers produce comparable nectar volumes regardless of the time of day they open. This pattern contrasts with the circadian fluctuations in nectar production reported in other plant species, where nectar secretion peaks coincide with the activity periods of primary pollinator guilds (Matile 2006, Luo et al. 2011, Edge et al. 2012, Prieto-Benítez et al. 2016, Wang et al. 2025). This finding suggests that apples do not rely on a single pollinator guild and instead attract insect visitors throughout the 24-h cycle and that nocturnal visitation by insects is not happenstance.

Visitation alone does not guarantee pollination (Joshi et al. 2020 July 10), and distinguishing true pollinators from incidental visitors is particularly challenging in nocturnal systems. Studying nocturnal pollinators remains methodologically difficult, especially when attempting to generate comparable abundance estimates across the full diel cycle (Wong and Didham 2024). In our study, insect sampling methods differed in their specificity: only the video recordings targeted flower visitation directly, whereas transects and light traps sampled broader nocturnal insect activity. Additionally, insufficient camera resolution prevented reliable species-level taxonomic identification of visitors. Together, these constraints underscore the need for improved nocturnal monitoring tools to better resolve the identity and effectiveness of nighttime pollinators in apple and other crops. Targeted pollen-load analyses of captured insects would help determine which taxa are actually transferring apple pollen, and metabarcoding of residual environmental DNA on stigmas or corollas could reveal the full suite of floral visitors, including those undetected by traditional sampling. Further evidence for adaptation of apple flowers to nighttime pollination would come from directly linking volatile release to insect responses, as well as from transcriptomic studies testing whether nocturnal visitation induces changes in floral gene expression. Integrating such approaches with standardized nocturnal observations would allow a more definitive separation of effective pollinators from background insect activity, sharpening our understanding of apple’s diel pollination cycle.

## 5. Conclusion

Taken together, our results demonstrate that apple flowers function as around-the-clock pollinator resources, supporting both diurnal and nocturnal insects through stable nectar rewards and temporal changes in floral volatile composition. Nectar availability remained consistent across the 24-h cycle, while volatile bouquets extracted from crushed flowers shifted dielly and interannually in ways consistent with the attraction of night-active dipterans and moths alongside daytime bee pollinators. These patterns, combined with direct evidence of nocturnal flower visitation, indicate that nighttime insect activity is not incidental but may represent a meaningful component of apple pollination ecology. From a practical perspective, recognizing nocturnal insects as contributors to pollination has direct implications for orchard management: pest control strategies and pesticide application guidelines developed with an exclusively diurnal focus risk underestimating impacts on beneficial nocturnal pollinators. These findings underscore the importance of considering pollinator activity across the entire diel cycle when developing orchard management practices, including decisions about pesticide use during bloom.

## 6. Data Availability

All numerical data used in analyses, as well as exemplary videos of nocturnal pollination, are available from the Dryad Digital Repository (https://datadryad.org/share/LINK_NOT_FOR_PUBLICATION/jlEo5-37XA0OusYENMpFZM_WxbG1STQPdFfQmPrI0qc). Original video footage is available from the corresponding author upon request.

## 7. Acknowledgements

This research was funded by the University of Arkansas through a Chancellor Innovation Award to ELW and NJ as well as Honors College Research Grants to MJ, CM, and EB, and by NSF IOS-2238931 to ELW. We thank Erik Pollock for his help with GC-MS volatile analysis, Chance Powell for his help with 24-hour video recordings, Kierstan Taylor and Matthew Brownshadel for their help with video analysis, and the Milo J. Shult Agricultural Research and Extension Center for access to the experimental apple orchard. We also thank anonymous reviewers for their comments on early drafts of the manuscript.

## Conflicts of Interest

The authors declare that they have no known competing financial interests or personal relationships that could have appeared to influence the work reported in this paper.

## Author Contributions

Conceptualization and Methodology: SP, DK, NJ, ELW. Investigation: SP, DK, MJ, EB, CM, MTL, GH, NTP, NJ, ELW. Data curation and Formal analysis: SP, DK, MJ, EB, CM, TH, ELW. Visualization: DK, SP. Funding acquisition: MJ, CM, EB, NJ, ELW. Project administration: SP, NJ, ELW. Supervision: NJ, ELW. Writing – original draft: DK, SP, MJ, EB, CM, ELW. Writing – review and editing: all authors.

## Notes

https://datadryad.org/share/LINK_NOT_FOR_PUBLICATION/jlEo5-37XA0OusYENMpFZM_WxbG1STQPdFfQmPrI0qc

### Competing Interest Statement

The authors have declared no competing interest.

