## Supplementary Fig. for "Apple floral attractants function across the 24-hour cycle to support potential diurnal and nocturnal pollinators"

**SUPPLEMENTARY FIGURES**

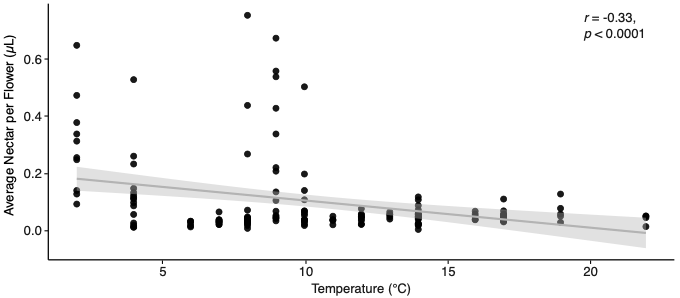

Fig. S1. Relationship between floral nectar volume and air temperature. Each point represents the mean nectar volume per tree (N = 156). The fitted line, with a 95% confidence interval, and reported statistics are based on Pearson’s correlation and are shown for illustrative purposes only. See the main text for results from the linear mixed-effects model.

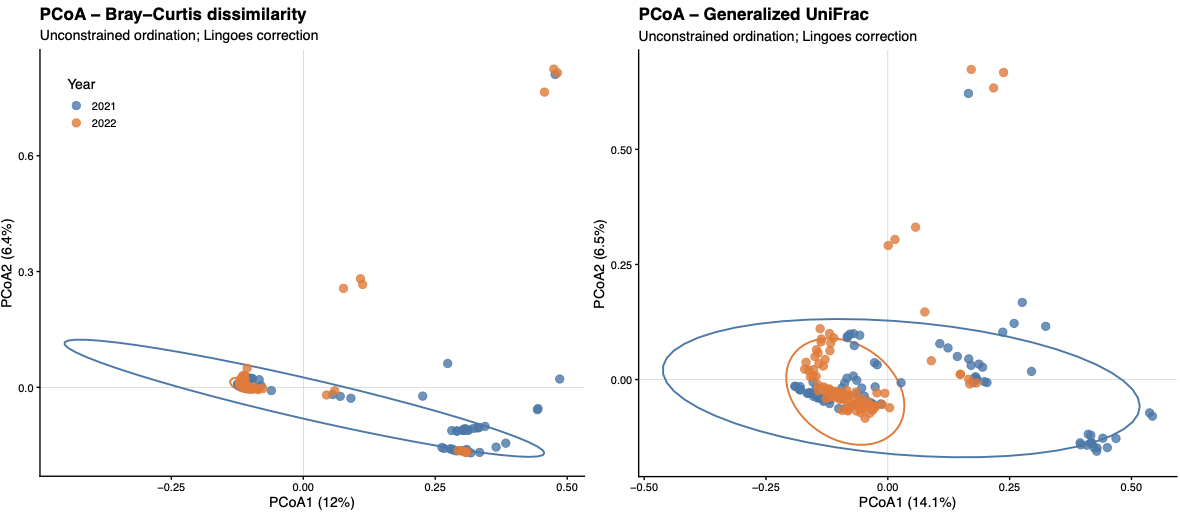

Fig. S2. Floral volatile composition differed strongly between years in both Bray-Curtis dissimilarity (differences in compound identity and relative abundance) and Generalized UniFrac dissimilarity (relative abundance and structural dissimilarity). When entered first in the sequential PERMANOVA, the effect of year was significant for both dissimilarity measures (Bray-Curtis: pseudo-*F* = 14.4, *P* = 0.0001; Generalized UniFrac: pseudo-*F* = 11.9, *P* = 0.0001) and remained significant in marginal PERMANOVA after accounting for the effects of time of day and weather variables (Bray-Curtis: pseudo-*F* = 15.3, *P* = 0.0001; Generalized UniFrac: pseudo-*F* = 12.8, *P* = 0.0001; df = 1, 176). Permutation tests for homogeneity of multivariate dispersions were also highly significant (*P* = 0.0001), indicating that the years differed in heterogeneity of volatile composition, not just in their mean composition. Differences between 2021 and 2022 are visualized using principal coordinates analysis (PCoA). Points represent individual samples; ellipses indicate 95% confidence regions for each year.

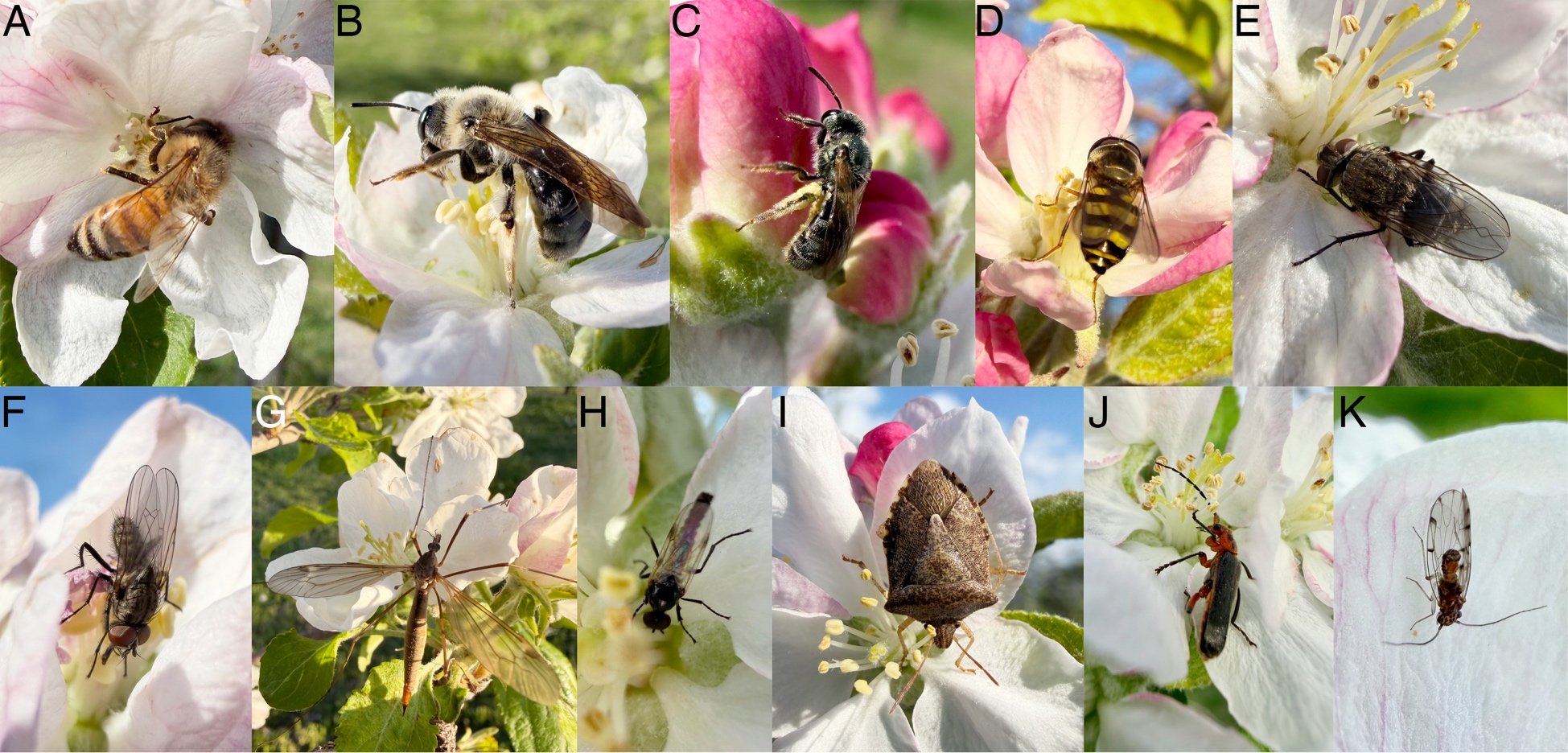

Fig. S3. Insects visiting apple flowers during the day in Northwest Arkansas, not to scale: (A) honeybee *Apis mellifera*; (B) mining bee *Andrena* sp. (C) sweat bee (subtribe Halictina); (D) hoverfly *Eupeodes* aff. *americanus* (Brachycera); (E, F) unidentified flies (Brachycera); (G) cranefly *Tipula* (*Triplicitipula*) sp. (Nematocera); (H) March fly *Dilophus* sp. (Nematocera); (I) shield bug *Euschistus servus*; (J) soldier beetle *Atalantycha bilineata*; (K) bark louse (Psocodea). Photos taken at the study site between March, 31 and April 9, 2026.

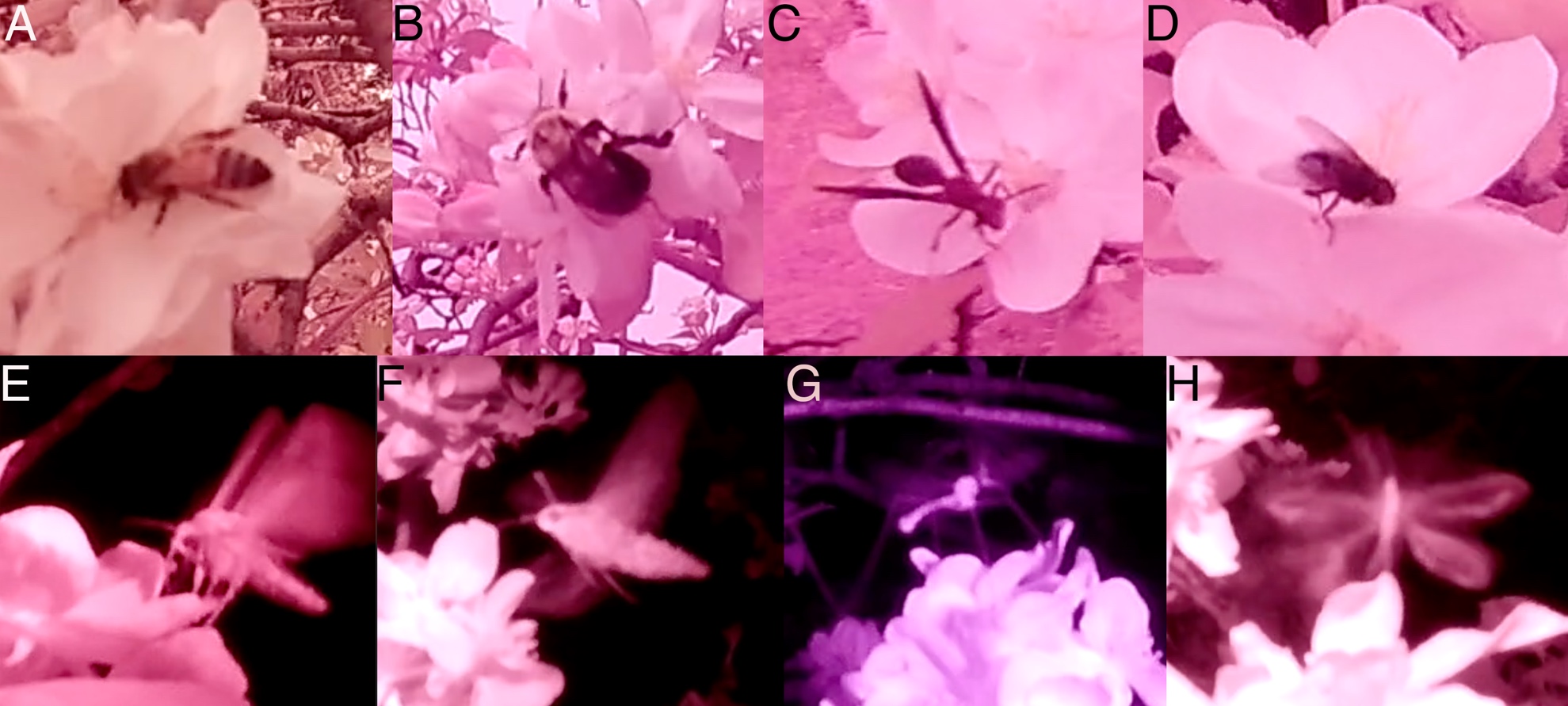

Fig. S4. Fragments of infrared footage showing various visitors of apple flowers during the day and at night, not to scale: (A) honeybee; (B) bumblebee; (C) paper wasp; (D) brachyceran fly; (E) noctuid moth; (F) hawkmoth; (G) cranefly (Nematocera); (H) lacewing.

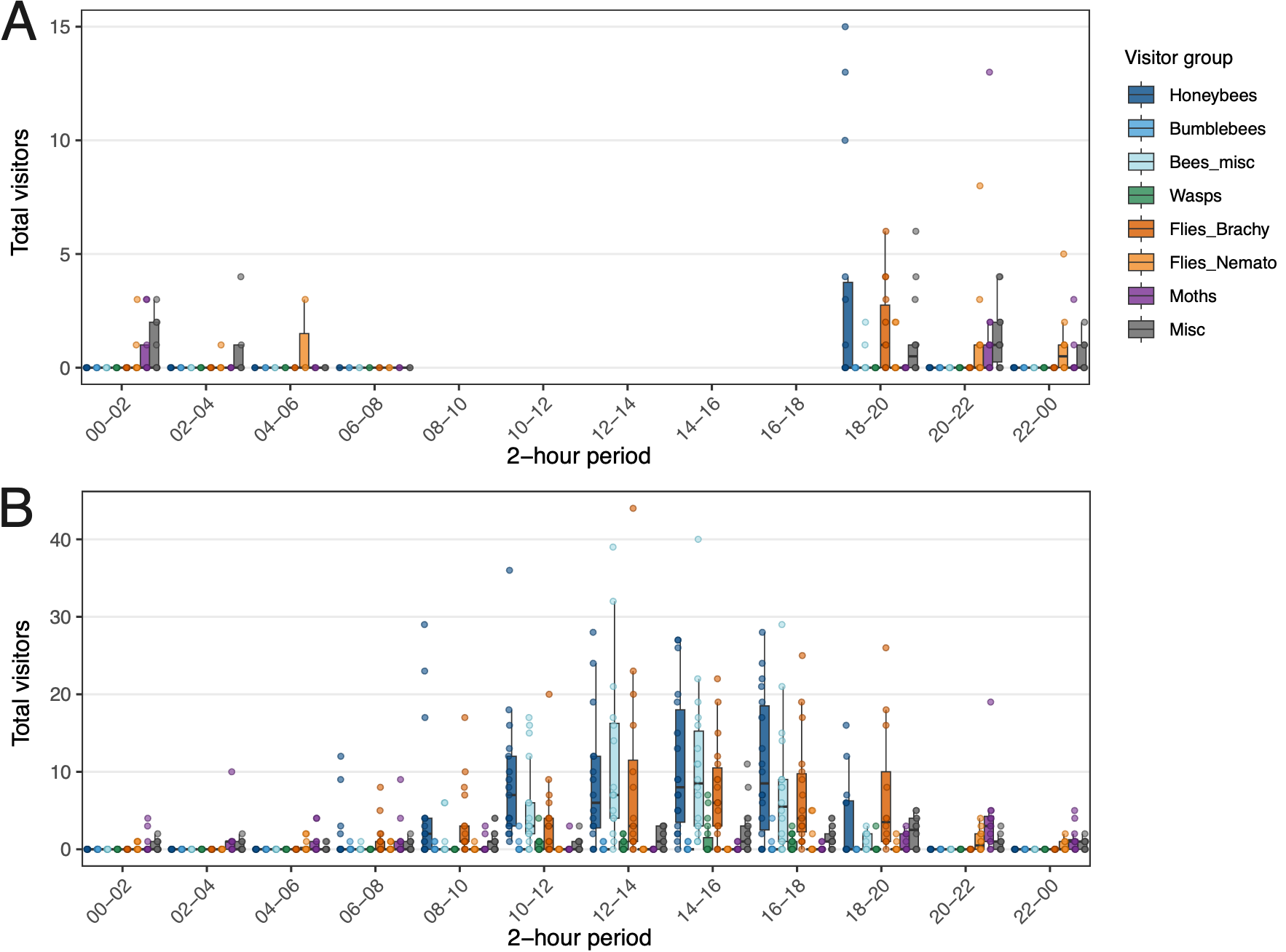

Fig. S5. Apple flower-visiting insects detected in video recordings in (A) 2021 and (B) 2026 (note different y-axes). Each point is an insect count for a given calendar date, flower cluster, and 2-h video period. Horizontal lines inside the boxes are medians (50th percentile), the height of the box is interquartile range (IQR), and whiskers extend to the most extreme observations within 1.5 × IQR.

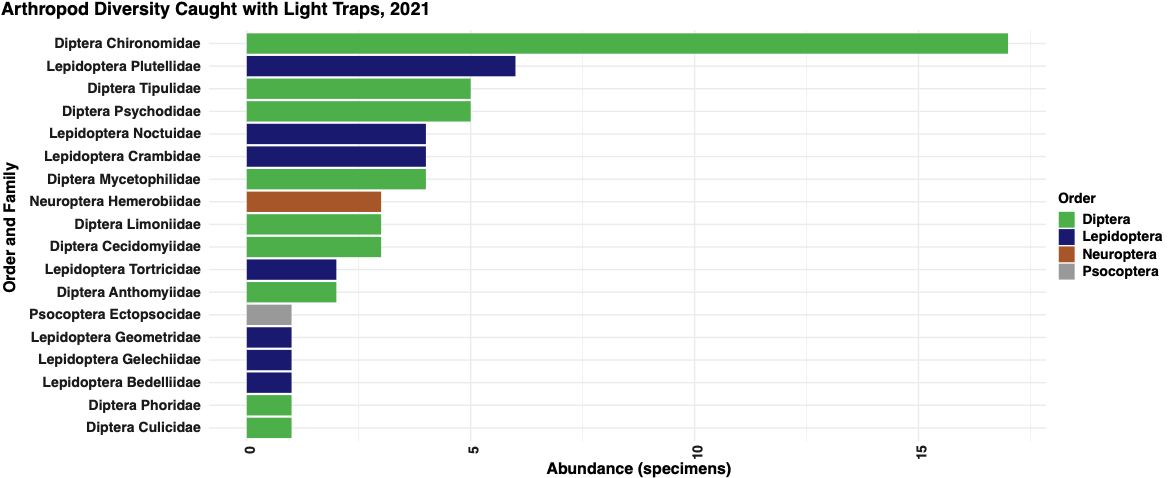

Fig. S6. The diversity of apple orchard-inhabiting nocturnal arthropods in light-trap samples in 2021.

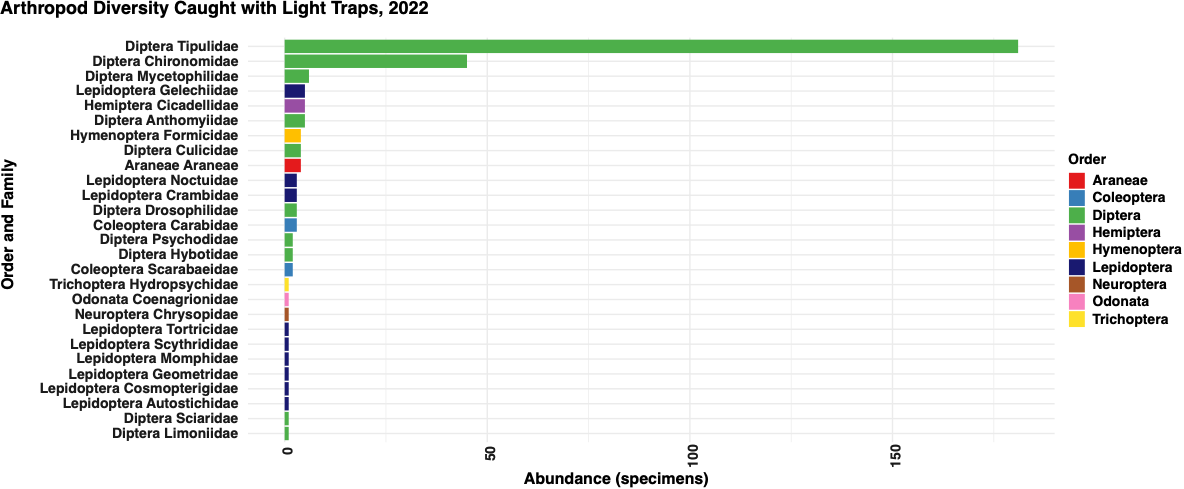

Fig. S7. The diversity of apple orchard-inhabiting nocturnal arthropods in light-trap samples in 2022.

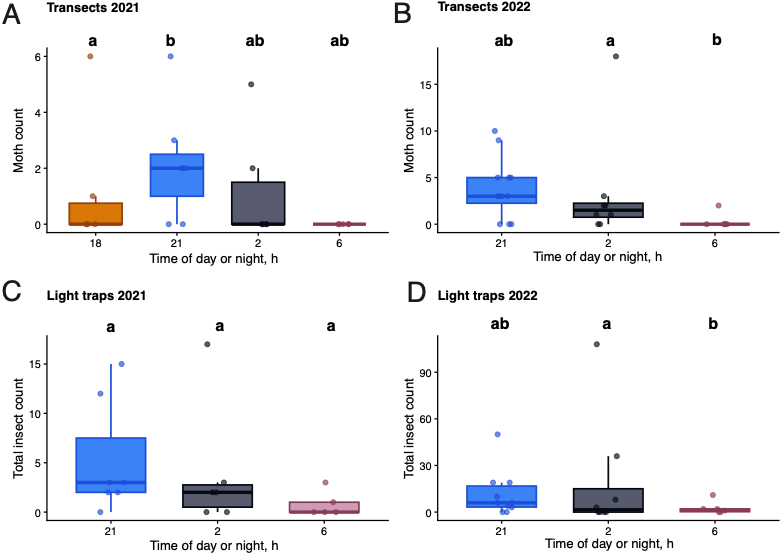

Fig. S8. Abundance of moths and other insects across the night. Data from transect surveys in (A) 2021 and (B) 2022. Data from light traps in (C) 2021 and (D) 2022. Horizontal lines inside the boxes are medians (50th percentile), the height of the box is the interquartile range (IQR), and whiskers extend to the most extreme observations within 1.5 × IQR. Different letters indicate significant differences among sampling times based on Tukey-adjusted pairwise comparisons (*P* < 0.05). Full GLMM statistics are provided in Supplementary Table S4.

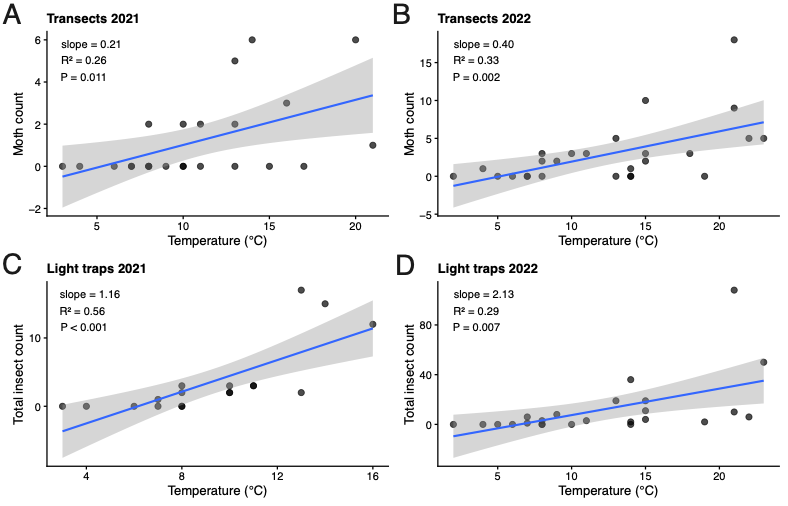

Fig. S9. Relationships between ambient temperature at night and insect abundance per sample in (A, B) transects and (C, D) light traps in 2021–2022. Lines represent descriptive linear regressions and are shown with 95% confidence intervals; these regressions are provided for visualization only. Full GLMM statistics are provided in Supplementary Table S4.
